## Supplementary for "A modeling framework for determining modulation of neural-level tuning from non-invasive human fMRI data"

### Supplementary Methods

#### Bayesian Estimation of Parametric Model

We developed two different hierarchical Bayesian estimation models, which differ in the number of hierarchical layers. Both versions estimated voxel-specific parameters, but the second version additionally estimated systematic variability across participants. A schematic of the first version, which collapsed across participants, is given by Supplementary Figure 1. As displayed in Supplementary Figure 1, every voxel was allowed its own value for preferred orientation ($\phi_v$), concentration parameter ($\kappa_v$), additive constant ($\alpha_v$), multiplicative constant ($\gamma_v$), and noise parameter ($\sigma_v$). In addition, for the high-contrast presentation (Equations 4a and 4b), rather than a low-contrast presentation (Equation 3), each voxel had its own modulation magnitude parameter, whether that occurred through an additive shift ($a_v$) or multiplicative gain ($g_v$). Because higher levels of contrast increase neural activity, the additive modulation parameter was constrained to be positive and the multiplicative modulation parameter was constrained to be greater than one.

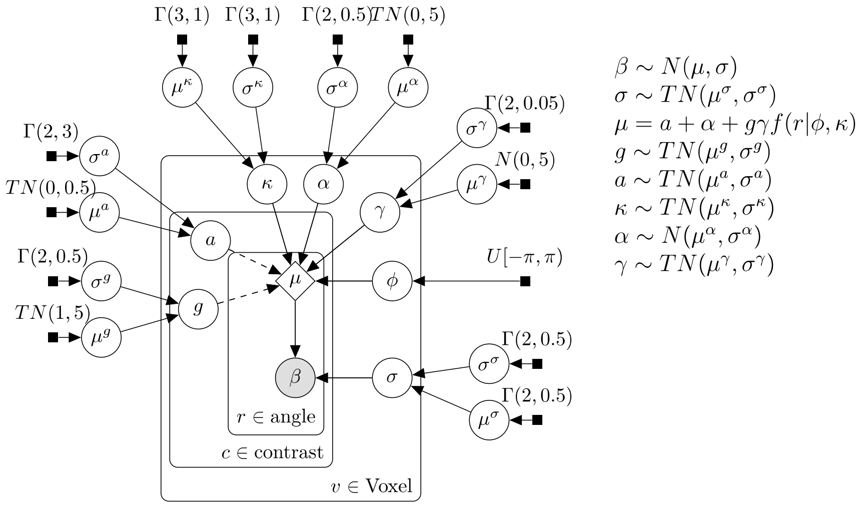


**Supplementary Figure 1** Schematic for Bayesian estimation of the parametric model, collapsing across participants. Filled square nodes indicate priors, open circles are estimated parameters, the shaded circle is the observed data, and the open diamond is the result of a deterministic function of the parameters (Equations 3 and 4). Nodes are grouped with the square "plates", indicating over which subsets of the data the node is replicated. The distribution assigned to each node is listed to the right of the diagram. $N(\mu,\sigma)$ is a normal with location $\mu$ and scale $\sigma$, and $TN(\mu,\sigma)$ is a normal with the same parameters, truncated below at $\mu$. $\Gamma(\zeta,\tau)$ is a gamma distribution with shape $\zeta$ and rate $\tau$. Parameters $\gamma$, $\kappa$, $a$, and $g$ follow truncated normal distributions, and $\alpha$ follows a normal distribution. Both kinds of modulation ($a$ and $g$) are depicted in this single diagram, but only one modulation was allowed during model fitting. In another version of the model, each of the $\mu^x$ parameters are themselves estimated hierarchically across participants.

The second, more complex version acknowledged differences in how participants responded to the contrast manipulation: e.g., perhaps one participant did not attend to the stimuli and so the influence of contrast was subject to floor effects. In this case, the participant's voxels would have, on average, smaller values for the modulation parameters ($a_v$ and $g_v$), relative to a more vigilant participant. Participant-level parameters in this second version were themselves analyzed hierarchically (i.e., there would be an additional plate in Supplementary Figure 1, covering participants). Both versions of the model led to the same conclusions in the collected dataset: a difference in favor of the multiplicative model, of 13.2 standard errors for the simpler version, and 14.5 for the more complex version, in units of expected log pointwise-predictive density, a measure of the predictive ability of a model[4].

Model recovery simulations were run with the more complex version. In the recovery simulations only, all voxels shared a single noise parameter. Estimating only a single noise parameter did not alter the main conclusions of the model comparison when it was applied to real data; the multiplicative model was still preferred. We report the version with voxel-specific noise since this version was preferred in model comparison on the real data (for both forms of modulation) against the version with a fixed noise term (see also Supplementary Figure 1).

#### Non-Parametric Check

***Multiplicative and Additive modulation can be distinguished by slope of activation plots.***

In the main text, we suggested that additive versus multiplicative modulation could be assessed by looking at the slope of the neural tuning function plotted at high contrast (the modulated condition) against the same tuning function plotted at low contrast (the baseline condition). Here, we show why. For this proof, we assume a discrete number of neurons, $N_v$, that contribute to the fMRI BOLD response of a particular voxel (each voxel can take on a different integer $N_v$). In contrast to Equation 2 in the main text, we start with a more general expression of the Neural Tuning Function, $NTF_i(\cdot)$,of a particular neuron, $i$, evaluated at a particular value of the stimulus dimension, $r$, (e.g., that neuron's response when presented with a grating of orientation $r$).

$$NTF_i(r) = \alpha_i + \gamma_i f_i(r) \tag{1}$$

Note that this NTF is presented in its most general form such that every neuron could have a differently shaped tuning function, $f_i$, including the possibility of multimodal tuning. In addition, every neuron could have a unique additive baseline response, $\alpha_i$, and unique multiplicative constant, $\gamma_i$. If the NTF of Supplementary Equation 1 indicates the metabolic cost of each neuron's activity that contributes to the BOLD signal, rather than literal firing rate, then the voxel response to stimulus, $r$, is simply the sum of the NTFs that contribute to the voxel, as shown in Supplementary Equation 2, which replaces the sum of the additive baselines responses with a voxel specific additive term, $\alpha_v$.

$$\begin{aligned} VTF_{baseline}(r) &= \sum_{i=1}^{N_v} \{\alpha_i + \gamma_i f_i(r)\} \\ &= \sum_{i=1}^{N_v} \alpha_i + \sum_{i=1}^{N_v} \gamma_i f_i(r) \\ &= \alpha_v + \sum_{i=1}^{N_v} \gamma_i f_i(r) \end{aligned} \tag{2}$$

This equation captures the voxel response in some baseline condition. Next, we implement multiplicative modulation by assuming that all neurons that contribute to the voxel experience the same magnitude of multiplicative modulation, $g_v$, similar to Equation 4a in the main text, resulting in Supplementary Equation 3. This equation rearranges the terms with the goal of representing the multiplicative VTF in terms of the baseline VTF, to specify the regression equation when plotting voxels response in the modulated condition as a function of response in the baseline condition:

$$\begin{aligned} VTF_{multiplicative}(r) &= \sum_{i=1}^{N_v}\{\alpha_i + g_v\gamma_i f_i(r)\} \\ &= \alpha_v + g_v \sum_{i=1}^{N_v} \gamma_i f_i(r) \\ &= g_v \left\{ {\alpha_v}/{g_v} - \alpha_v + \left[\alpha_v + \sum_{i=1}^{N_v} \gamma_i f_i(r)\right]\right\} \\ &= g_v\{{\alpha_v}/{g_v} - \alpha_v + VTF_{baseline}(r)\} \\ &= \{\alpha_v - g_v\alpha_v\} + g_v VTF_{baseline}(r) \end{aligned} \tag{3}$$

In other words, the predicted voxel response in the modulated condition (e.g., high contrast, rather than low contrast) should be the multiplicative modulation constant times the voxel response in the baseline condition plus an intercept that reflects both the multiplicative modulation constant and the voxel specific additive term. This is true regardless of the tested stimulus, $r$. Thus, the slope of the orthogonal regression relating the modulated condition to the baseline condition should be $g_v$ for this particular voxel. Each voxel could have a different multiplicative modulation constant, and thus a different slope, but if the modulated condition tends to produce a larger BOLD response, then the multiplicative modulation constants should tend to be greater than 1.0 on average (i.e., average slope > 1).

For additive modulation, it is assumed that all neurons contributing to the voxel experience the same magnitude of additive modulation, $a_v$, similar to Equation 4b in the main text, resulting in Supplementary Equation 4:

$$\begin{aligned} VTF_{additive}(r) &= \sum_{i=1}^{N_v}\{a_v + \alpha_i + \gamma_i f_i(r)\} \\ &= N_v a_v + \left[\alpha_v + \sum_{i=1}^{N_v} \gamma_i f_i(r)\right] \\ &= N_v a_v + VTF_{baseline}(r) \end{aligned} \tag{4}$$

In other words, the predicted voxel response in the modulated condition should be the additive modulation constant (times the number of neurons contributing to the voxel) added to the response in the baseline condition. This is true regardless of the tested stimulus, $r$. Thus, the orthogonal regression relating the modulated condition to the baseline condition should have an intercept of $N_v a_v$ and a slope exactly equal to 1.0. Each voxel could have a different additive modulation, and thus a different intercept, but if the modulated condition tends to produce a larger BOLD response, then the values of the additive modulation should tend to be greater than zero on average (i.e., average intercept > 0), but the slopes of all voxels should be 1.0.

This proof makes no assumptions about the shapes of the neural tuning functions, and it makes no assumptions that the neurons contributing to each voxel have the same shape. Each neuron is allowed to have its own unique tuning function. The one key assumption made in the slope test (an assumption that is shared with the parametric model), is that all neurons contributing to a particular voxel have the same multiplicative modulation or the same additive modulation. However, we can consider relaxations of this assumption, in particular to the less constraining (and fairly plausible) assumption that if the magnitude of the multiplicative or additive modulation varies across neurons within a voxel, it does not vary systematically with the orientation preference of the neurons. If the modulation constant varies systematically across neurons contributing to a voxel, then it is possible that multiplicative modulation could produce a slope of 1.0 and that additive modulation could produce a slope greater than 1.0. For instance, if the additive modulation was greater for the neurons that preferred stimuli that were also preferred by the voxel, additive modulation could produce a slope greater than 1.0. Analogously, if the multiplicative modulation was smaller for the neurons that preferred stimuli that were also preferred by the voxel, multiplicative modulation could produce a slope of 1.0. Such confounding relationships between neural preferences and voxel preferences may occur by chance for some voxels, but there is no obvious reason to expect such a systematically confounding relationship to occur for most voxels. Thus, it is likely that this assumption could be relaxed, and provided that heterogeneity of modulation magnitude is uniformly applied across the neurons contributing to a voxel, the outcome of the slope test should be reliable.

***Bayesian estimation of the non-parametric test.***

When the noise differs at high and low contrast, Equation 5 in the main text will produce a biased estimate of the slope. The contrast of a stimulus can affect noise[5], an effect that was visible in our data (Supplementary Figure 2). Given this potential for a biased slope, we supplemented the frequentist non-parametric test with Bayesian estimation of the slope that did not assume equal noise.

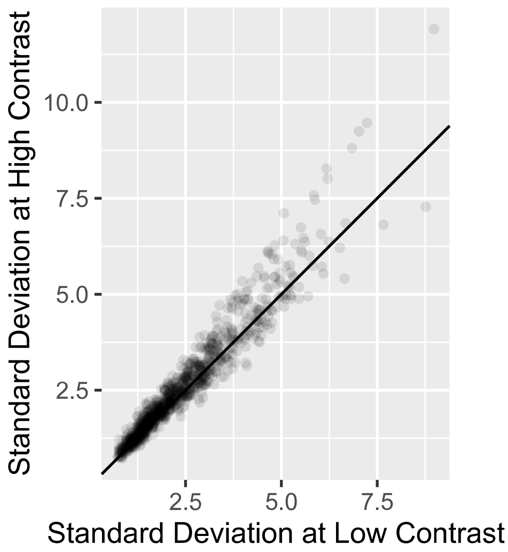


**Supplementary Figure 2** Within voxels, the noise at high contrast is slightly higher than noise at low contrast. Each point represents the pooled standard deviation for a voxel at high and low contrast (i.e., the variability of a voxel's beta, pooled across orientations). The solid line marks the diagonal.

As before, let $x$ be the activity of a voxel at low contrast. Additionally, let $j$ be an index for run and $r$ be an index for orientation. The low-contrast activity is assumed to vary around location parameters that depend on orientation, $\zeta_r^x$, with the standard deviation given by $\sigma_x$ (i.e., noise at low contrast).

$$x_{r,j} \sim N(\zeta_r^x, \sigma^x) \tag{5}$$

Supplementary Equation 5 defines the likelihood function for voxel activity at low contrast. The likelihood for high contrast activity, $y$, is analogous:

$$y_{r,j} \sim N(\zeta_r^y, \sigma^y) \tag{6}$$

The location parameters, $\zeta_r^x$ and $\zeta_r^y$, in Supplementary Equations 5 and 6 are given by deterministic functions of the output of voxel tuning function at low contrast. We denote that output with $z_r$. The $\zeta_r^x$ are equal to $z_r$, while the $\zeta_r^y$ are shifted and scaled according to an offset, $a$, and gain, $g$.

$$\begin{aligned} \zeta_r^x &= z_r \\ \zeta_r^y &= a + g z_r \end{aligned} \tag{7}$$

In the case of only additive modulation, the offset $a$ in Supplementary Equation 7 gives the magnitude of the modulation (i.e., the $a = N_v a_v$ according to Supplementary Equation 4). In the case of only multiplicative modulation, the offset $a$ will be a function of both the baseline offset and the multiplicative modulation (i.e., $a = \alpha_v - g_v \alpha_v$ according to Supplementary Equation 3). The variable $g$ is the same slope as estimated by orthogonal regression. But in contrast to the estimate provided by Equation 5 in the main text, this Bayesian model allows that the noise at high contrast, $\sigma^y$, may differ from the noise at low contrast, $\sigma^x$, mitigating this source of bias in the orthogonal regression estimate of slope.

Supplementary Equations 5 – 7 resemble the Bayesian model presented in the main paper. However, while that model defined a parametric equation for the voxel tuning function, here we simply model it with a normal distribution centered around a mean $\mu^z$ with standard deviation $\sigma^z$.

$$z_r \sim N(\mu^z, \sigma^z) \tag{8}$$

We estimated this model hierarchically, allowing $\mu^z$, $\sigma^z$, $z_r$, $a$, $g$, $\sigma^x$ and $\sigma^y$ to vary by voxel (Supplementary Figure 3).

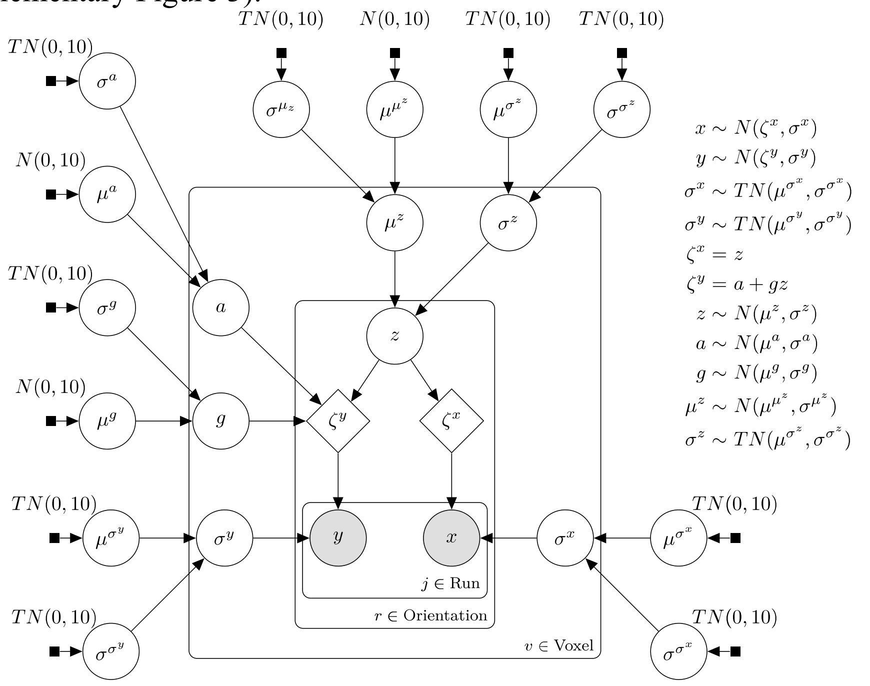


**Supplementary Figure 3** Schematic of the Bayesian implementation of the non-parametric check (Supplementary Equations 8-11). See also caption of Supplementary Figure 1.

For this Bayesian model, we sampled 1000 draws from the estimated posterior distribution in each of five chains, following 1000 warmup draws per chain. The resulting posterior distributions suggest that the noise at high contrast was slightly higher than the noise at low contrast (Supplementary Figure 4A). However, supporting the orthogonal regression estimate, the posterior distribution of the average slope was above 1 (Supplementary Figure 4B).

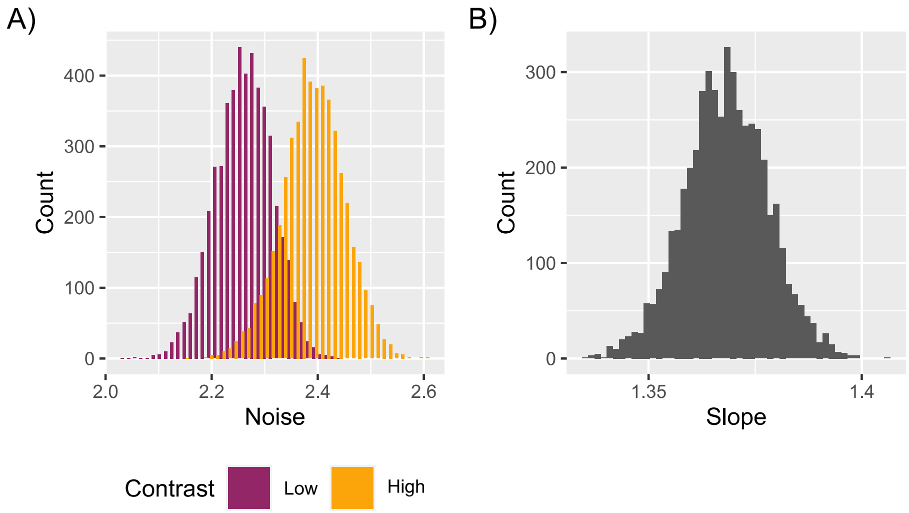


**Supplementary Figure 4** Posterior distributions from the Bayesian estimation of the non-parametric check. **A)** Voxels' activity at high contrast may be noisier than their activity at low contrast. The histograms give the posterior samples for the location parameters of the population of distribution (across voxels) of the noise at low, $\mu^{\sigma^x}$, and high, $\mu^{\sigma^y}$, contrast. **B)** The posterior samples for the average (across voxels) slope, $\mu^g$, are above 1.

#### Supplementary Results

##### Behavior

Average accuracy for the spatial frequency change detection task was 72% and 74% for low and high-contrast gratings, respectively ($p = 0.13$). Eye-tracking was used to assess whether participants maintained adequate fixation in the scanner: over 90% of all participants' fixations ended within 2° of the location of the run's average fixation (range: 93–99%).

##### Population Receptive Field Mapping

Only voxels from V1 were analyzed, and only voxels whose population receptive fields overlapped with the stimulus (Methods). This resulted in 1,010 voxels, ranging from 90 to 188 across participants.

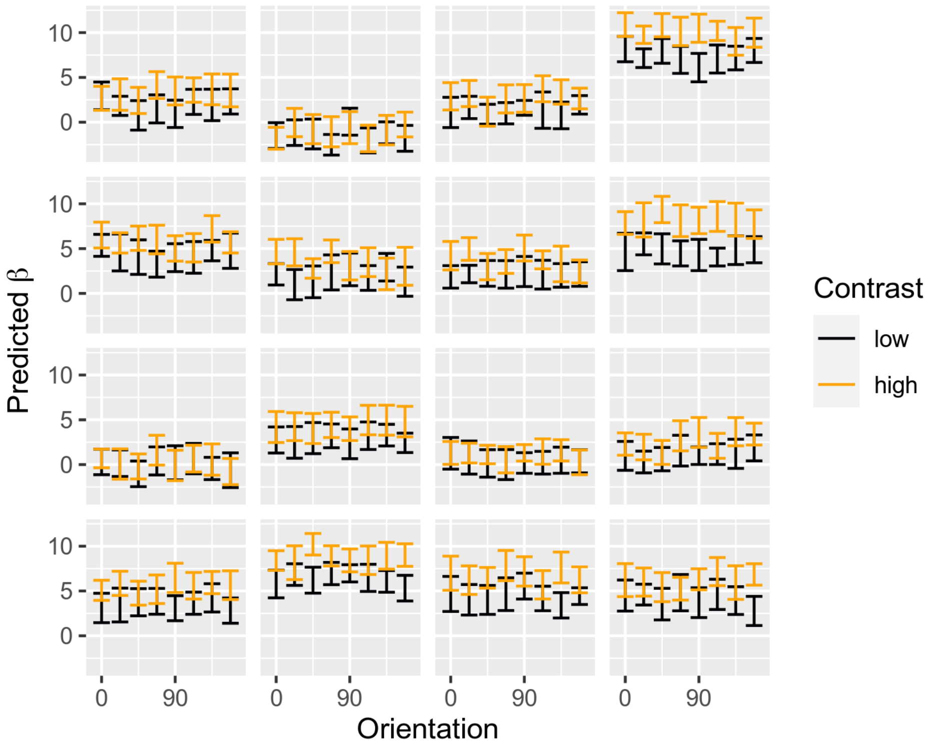


**Supplementary Figure 5** Samples from the Posterior Predictive Distribution of the Multiplicative Model. Data plotted as in Figure 4B from the main text. Simulated voxels capture the qualitative trends of real voxels (i.e., only minor tuning across orientation, low signal-to-noise, higher activity with high as compared to low contrast).

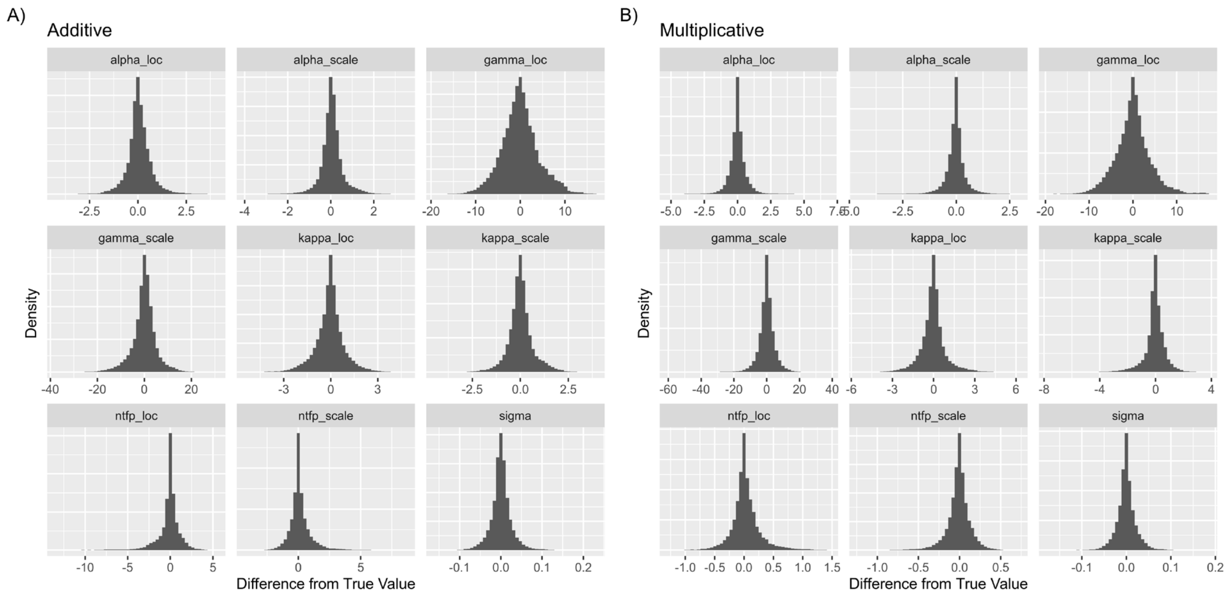


**Supplementary Figure 6** Parameter Recovery for Data-Informed Model Recovery. Distributions show the difference from true value across draws when **(A)** the additive model was fit to datasets generated from the additive posterior predictive distribution, or **(B)** the multiplicative model was fit to datasets generated from the multiplicative posterior predictive distribution. In all cases, the distributions are centered on 0, indicating lack of bias in the estimated parameters. Panels give parameters (compare to Supplementary Figure 1): alpha_loc: $\mu^{\alpha}$; alpha_scale: $\sigma^{\alpha}$; gamma_loc: $\mu^{\gamma}$; gamma_scale: $\sigma^{\gamma}$; kappa_loc: $\mu^{\kappa}$; kappa_scale: $\sigma^{\kappa}$; ntfp_loc: either $\mu^{a}$ or $\mu^{g}$; ntfp_scale: either $\sigma^{a}$ or $\sigma^{g}$; sigma: $\mu^{\sigma}$.
